# Glucose concentration of neuronal media formulations influences PINK1-dependent mitophagy in human iNeurons

**DOI:** 10.1101/2025.04.01.646648

**Authors:** Benjamin O’Callaghan, Daniela Melandri, Darija Soltic, Katharina Cosker, Marc P.M. Soutar, James R. Evans, Hannah Lucas-Clarke, Joe Robin, Sonia Gandhi, Nicol Birsa, Charles Arber, Selina Wray, Helene Plun-Favreau

**Affiliations:** Department of Neurodegenerative Diseases, UCL Queen Square Institute of Neurology, London, UK; Aligning Science Across Parkinson’s (ASAP) Collaborative Research Network, Chevy Chase, MD, USA, 20815; UCL Alzheimer’s Research UK, Drug Discovery Institute, London, UK; The Francis Crick Institute, 1 Midland Road, London, UK; Department of Clinical and Movement Neurosciences, UCL Queen Square Institute of Neurology, London, UK; Department of Neuromuscular Diseases, UCL Queen Square Institute of Neurology, London, UK

**Author notes:** Correspondence to Benjamin O’Callaghan and Helene Plun-Favreau.

**Keywords:** iNeuron, Mitophagy, mitoSRAI, Parkin, PINK1, pUb(Ser65)

## Abstract

Parkinson’s disease associated proteins PINK1 and Parkin collaboratively regulate stress-induced mitophagy. While *in vitro* human neuronal cultures are valuable for studying the roles of PINK1 and Parkin in a disease-relevant context, the impact of culture conditions on these processes remains largely underexplored. Here, it is shown that human induced neurons (iNeurons) cultured in N2B27 and BrainPhys medium exhibit distinct PINK1-Parkin dependent mitophagy phenotypes. Specifically, BrainPhys-cultured iNeurons show greater resistance to PINK1-dependent mitophagy initiation, linked to a reduction in glucose availability and reduced PINK1 protein availabilities, leading to decreases in stress-induced and basal mitophagy fluxes. These findings highlight the critical impact of culture conditions on mitophagy dynamics and emphasise the need to account for media-specific differences when using *in vitro* models to investigate mitophagy mechanisms in human neurons.

## INTRODUCTION

PTEN-induced kinase 1 (PINK1) and Parkin E3-ubiquitin ligase act in a common cellular pathway responsible for selective clearance of damaged mitochondria through autophagy (mitophagy) which is crucial for maintenance of a healthy mitochondrial network [1]. The identification of loss of function mutations in the genes encoding PINK1 [2] and Parkin [3] (*PINK1* and *PRKN* respectively) in individuals with early-onset and autosomal-recessive forms of the neurodegenerative disease Parkinson’s disease (PD) has underscored the potential importance of mitophagy in human neurons, in particular dopaminergic neurons of the substantia nigra.

A significant portion of our understanding of the PINK1-Parkin dependent mitophagy process has been derived from functional studies conducted in cancer cell line models, often involving the exogenous overexpression of Parkin and/or PINK1[4]. This work has provided great insight into the underlying mechanisms by which PINK1-Parkin orchestrate the selective tagging and clearance of damaged mitochondria for autophagosome-lysosome clearance. Under basal conditions, healthy mitochondrial units maintain a polarised mitochondrial membrane which facilitates the efficient partial import of PINK1’s N-terminus and its subsequent cleavage by the inner mitochondrial membrane resident Presenilin-associated rhomboid-like protease (PARL), rendering it vulnerable to cytoplasmic proteasomal degradation [5,6]. Upon mitochondrial damage and associated mitochondrial membrane potential (MMP) depolarisation, PINK1 is instead stabilised on the outer mitochondrial membrane (OMM) [7–9] where it activates through trans-autophosphorylation [10]. Activated PINK1 phosphorylates ubiquitin molecules in proximity of the OMM at Ser65 (pUb(Ser65)) which serve as adapter sites for the localised recruitment and activation of Parkin [11–14]. Full activation of Parkin also necessitates direct phosphorylation by PINK1, also at Ser65 of a ubiquitin-like domain; pParkin(Ser65) [15]. Fully activated pParkin(Ser65) catalyses the deposition of further ubiquitin molecules on OMM proteins, which are then also phosphorylated by PINK1 leading to a positive feedforward cycle which efficiently coats damaged mitochondrial units with pUb(Ser65) [16]. These pUb(Ser65) tags serve as recognition sites for autophagy adapters such as NDP52 and optineurin, triggering autophagosome biogenesis, engulfment of the mitochondria and downstream lysosomal-dependant degradation [17–19].

At first it was unclear, whether the major steps of the PINK-Parkin mitophagy process described in proliferative cell line models also apply to neurons with endogenous Parkin expression levels [20–22]. Since, evidence has accumulated showing that PINK1 recruits and activates Parkin at damaged mitochondria of neurons, leading to deposition of pUb(Ser65) and subsequent recruitment of the autophagosome machinery [23–38]. This includes work by our group [35] and others [33,37,38] showing that PINK1-Parkin dependent activity and downstream mitophagy is readily detectable in iNeurons differentiated from human pluripotent stem cells (hPSCs) via overexpression of neurogenin-2 [33,35,37,38]. Despite these advancements, there remains large discrepancies in specific observations between studies and neuron models. Some of these are likely attributed to differences in species, neuronal subtypes and neuron/culture ages utilised, in addition to the strategies employed for initiating PINK1-Parkin mitophagy (e.g. specific chemical stressors to trigger mitochondrial depolarisation and/or oxidative stress). Another important related aspect that remains largely overlooked is that of culture medium composition.

It is well appreciated that cell culture conditions employed for *in vitro* neurological disease-modelling are far removed from the physiological *in vivo* settings of brain cells in living organisms. While 2D monoculture systems are extremely useful for robust mechanistic studies, cells are not exposed to the same complex biological architectures observed in the tissues of living organisms. These include contact-dependant cell-cell interactions and localised exposure to soluble factors released by neighbouring cells (e.g. substrates, growth factors and other signalling molecules). In the context of *in vitro* neuronal culture, base media compositions such as DMEM/F12 (a 1:1 mixture of Dulbecco’s Modified Eagle Medium (DMEM) and Ham’s F12) and Neurobasal (a modified version of DMEM/F12) supplemented with various vitamins, hormones, and antioxidants, often as part of neuronal supplement cocktails like N-2 or B27, have remained the most commonly used by the research community since their initial descriptions over three decades ago [39,40]. Research laboratories routinely utilise N2B27 medium, a 1:1 mix of N2-supplemented DMEM/F12 and B27-supplemented Neurobasal. These media have been essential for *in vitro* neurological disease modelling, however they were developed with a focus on supporting neuronal survival and/or growth rather than facilitating physiological neuronal functions. With that in mind, Bardy et al. developed a medium formulation to better support neurophysiological activity and which more closely recapitulates brain energy substrate availability [41].

Beginning with DMEM/F12 as the base, they showed that neuroactive amino acid concentrations could be reduced to facilitate both excitatory and inhibitory neurotransmission of hPSC-derived neurons. Together with modifications to several salt concentrations and a reduction in glucose to more physiological levels, this new base medium formulation named BrainPhys was shown to efficiently support hPSC-derived neuronal activity and long-term culture when supplemented with typical neuronal chemical supplements and growth factors.

In this study we demonstrate that neuronal medium composition is a key determinant of PINK1-Parkin-dependent mitophagy initiation and subsequent lysosomal delivery. hPSC-derived iNeurons cultured in BrainPhys showed attenuated deposition of pUb(Ser65) upon mitochondrial depolarisation, when compared to iNeurons cultured in N2B27, and this was largely linked to more limited glucose availability. This phenotype was at least partly explained by a post-transcriptional reduction in PINK1 protein availability. Using the mitoSRAI mitophagy sensor assay[42] in human neurons for the first time, the change in pUb(Ser65) deposition was correlated with a reduction in the downstream delivery of mitochondria to lysosomes in BrainPhys iNeurons. These results highlight the sensitivity of neuronal mitophagy to *in vitro* culture conditions and show the importance of culture medium selection when designing experiments focused on PINK1-Parkin dependent mitophagy.

## RESULTS

In order to compare the impact of using BrainPhys rather than N2B27 medium on neuronal PINK1-Parkin dependent mitophagy, iNeurons were first differentiated from a well characterised hPSC-line harbouring a TET-ON system for doxycycline-induced Ngn2 neuronal differentiation[43,44]. Following three days of doxycycline treatment, neural progenitor cells were then seeded in either BrainPhys or N2B27 (and glucose adjusted derivatives) and cultured for a further 21 days prior to experimental collection (Figure 1A, see methods and Table 1 for details on final media compositions).

**Figure 1.**
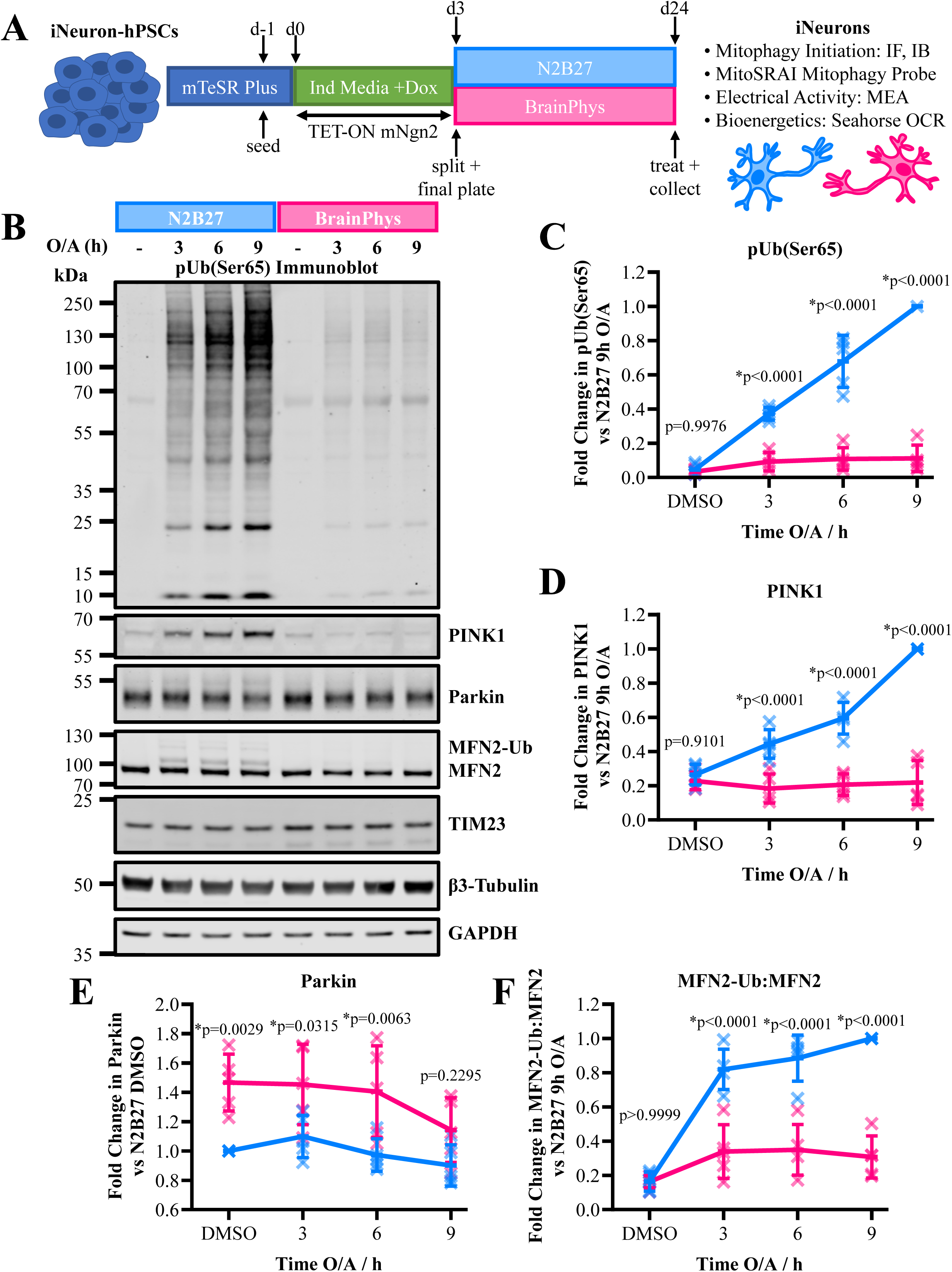
iNeurons cultured in BrainPhys medium are more resistant to PINK1-dependent mitophagy initiation. (**A**) Schematic of the iNeuron differentiation strategy and experimental workflow. (**B**) Representative immunoblots of d24 iNeurons cultured in N2B27 vs BrainPhys medium and treated with 1 µM O/A over a 9h time-course. (**C-F**) Quantification of pUb(Ser65) (**C**) PINK1 (**D**), Parkin (**E**) and the MFN2-monoUb:MFN2-FL ratio (**F**) from (**B**) (n=5 inductions, two-way ANOVA with Šidák *post hoc* correction). Data are shown as the mean with error bars□± SD from biological replicates, with normalised values for each biological replicate shown as individual markers.

### PINK1-dependent mitophagy initiation is attenuated by culturing iNeurons in BrainPhys over N2B27

PINK1-Parkin dependent mitophagy was triggered using a 1 µM equimolar mixture of oligomycin and antimycin-A (O/A) to halt the electron transport chain and depolarise the inner mitochondrial membrane (IMM). Immunoblot analysis of N2B27 iNeurons revealed a robust deposition of pUb(Ser65) over a 9h O/A time-course window (Figure 1B-C), in line with previous observations from iNeurons cultured in N2B27 [35] and Neurobasal-B27 [33,37]. In O/A-treated BrainPhys iNeurons, pUb(Ser65) deposition was significantly reduced (Figure 1B-1C). Immunofluorescence confocal imaging of N2B27 and BrainPhys iNeurons (Extended Data Figure 1A) corroborated these findings, with significant reductions in the intensity of O/A-induced pUb(Ser65) immunofluorescence within MAP2 positive neuronal regions also observed in BrainPhys vs N2B27 iNeurons (Extended Data Figure 1B). The reduction observed by confocal imaging was slightly subtler in magnitude. We hypothesise that this is most likely associated with the lower cell densities used in these imaging-based assays. Consistent with this, severe impairments in O/A-induced pUb(Ser65) deposition were also observed in hPSC-derived cortical neurons differentiated through dual SMADi approaches (Extended Data Figure 2A-B), which are typically cultured at much higher densities than iNeurons. Immunoblot revealed a marked reduction in PINK1 protein accumulation following O/A induced IMM depolarisation in BrainPhys iNeurons when compared to N2B27 iNeurons (Figure 1B and 1D). By contrast, Parkin protein levels were significantly elevated in BrainPhys iNeurons under basal conditions (Figure 1B and 1E). This is consistent with previous reports linking basal PINK1 protein availability to steady-state regulation of Parkin [31], most likely through background mitochondrial Parkin activation, subsequent autoubiquitination [45] and/or MARCH5-dependent degradation [46]. In line with reduced activation of Parkin by PINK1, ubiquitination of MFN2, a well described mitochondrial Parkin substrate was significantly attenuated in BrainPhys vs N2B27 iNeurons (Figure 1B and 1F). Overall, these data suggest that the reduction in PINK1 accumulation at least partly accounts for the decrease in pUb(Ser65) deposition and Parkin-dependent MFN2 ubiquitination.

### BrainPhys iNeuron pUb(Ser65) phenotype is mediated by non-transcriptional reduction in PINK1 availability

In order to determine whether the reduction in PINK1 protein accumulation and pUb(Ser65) deposition observed in BrainPhys iNeurons was transcriptionally driven, gene expression measurements through RT-qPCR were performed which revealed no discernible differences in *PINK1* mRNA levels between two media compositions (Figure 2A). Due to the dynamic nature of PINK1 protein regulation[47], we hypothesised that the non-transcriptional decrease observed in BrainPhys iNeurons could be due to reductions in protein synthesis/availability and/or destabilisation of interactions with the translocase of the outer membrane (TOM) complex. In order to distinguish between these potential contributing factors, iNeurons were treated with 10 µM MG132, a proteasome inhibitor. MG132 treatment allows detection of the mitochondrially imported and PARL cleaved PINK1 product which is otherwise degraded via the proteosome (Δ-PINK1; Figure 2B)[47]. This allowed the amount of PINK1 protein translated within a specific timeframe to be measured independently of mitochondrial depolarisation and TOM stabilisation. Similar to the reduction in full-length PINK1 (FL-PINK1) observed following 9h of O/A induced mitochondrial depolarisation, BrainPhys iNeurons also showed a reduction in levels of Δ-PINK1 stabilised by 9h of MG132 mediated proteasome inhibition (Figure 2C-2E). Co-application of O/A and MG132 (Extended Data Figure 3A-3C) further supports a global reduction in PINK1 protein availability as the underlying cause of reduced FL-PINK1 accumulation following O/A treatment. Notably, no overt change in the ratio between mitochondrial stabilised FL-PINK1 and Δ-PINK1 was observed following O/A and MG132 co-application, despite the overall reduction in PINK1 accumulation (Extended Data Figure 3A,3D). Importantly, MMP following 1µM O/A application is comparable between N2B27 and BrainPhys iNeurons.

**Figure 2.**
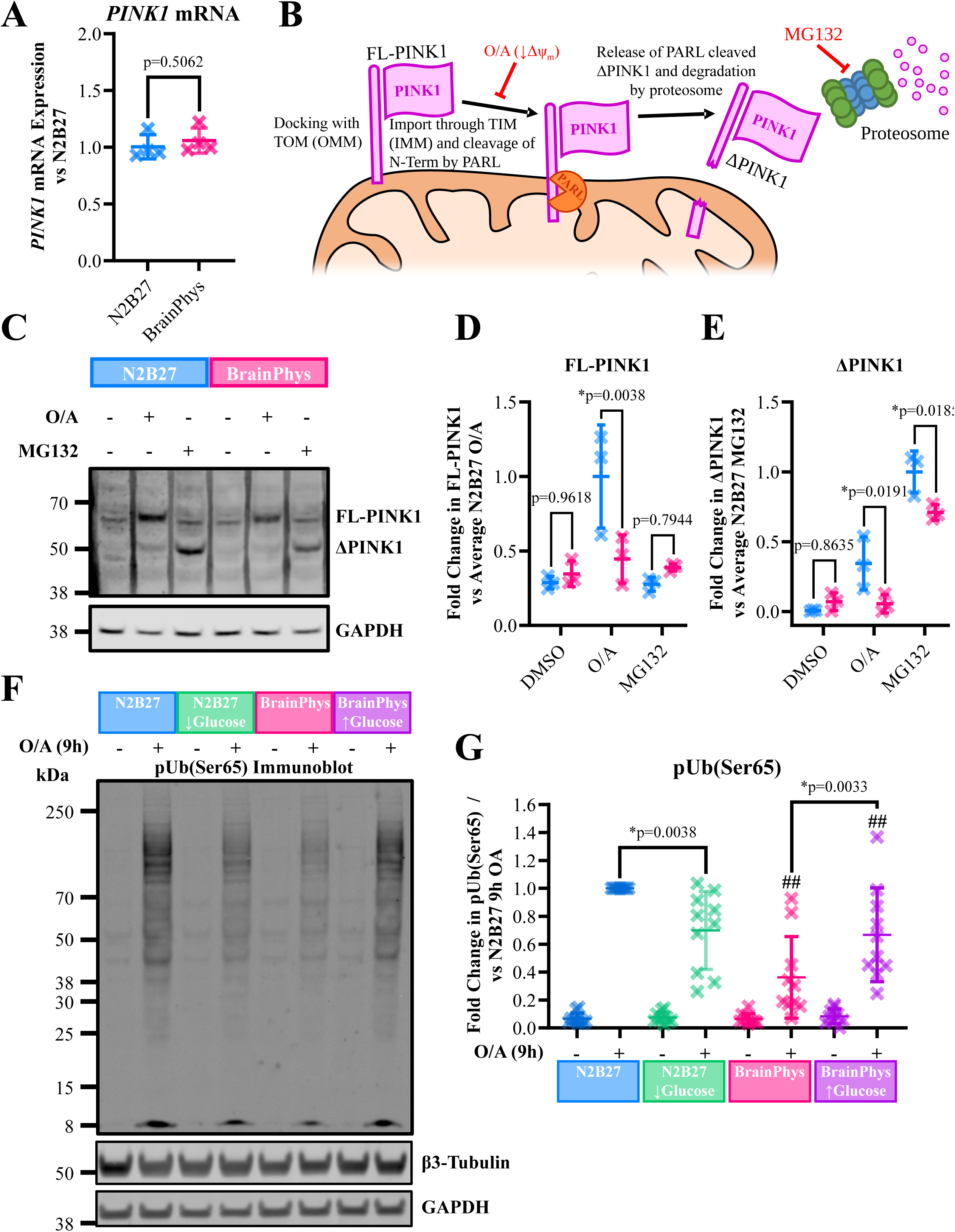
A post-transcriptional reduction in PINK1 protein availability and glucose driven differences in media composition contribute to pUb(Ser65) phenotype. (**A**) RT-qPCR quantification of *PINK1* mRNA in N2B27 and BrainPhys iNeurons (n=4 inductions, unpaired two-tailed t-test). (**B**) Representative immunoblots of d24 iNeurons cultured in N2B27 vs BrainPhys medium and treated with 1 µM O/A or 10 µM MG132 for 9h. (**C-D**) Quantification of full-length PINK1 (FL-PINK1) (**C**) and PARL-cleaved PINK1 (ΔPINK1) (**D**) from (**B**) (n=3 inductions, two-way ANOVA with Šidák *post hoc* correction). (**E**) Representative immunoblots of d24 iNeurons cultured in N2B27 vs BrainPhys medium with low 2.5mM glucose or high 21.25mM glucose and treated with 1 µM O/A for 9h. (**F**) Quantification of pUb(Ser65) from (**E**) (n=10 inductions, two-way ANOVA with Šidák post hoc correction, ## indicates p<0.005 vs N2B27 of same glucose concentration). Data are shown as the mean with error bars□±SD from biological replicates, with normalised values for each biological replicate shown as individual markers.

**Figure 3.**
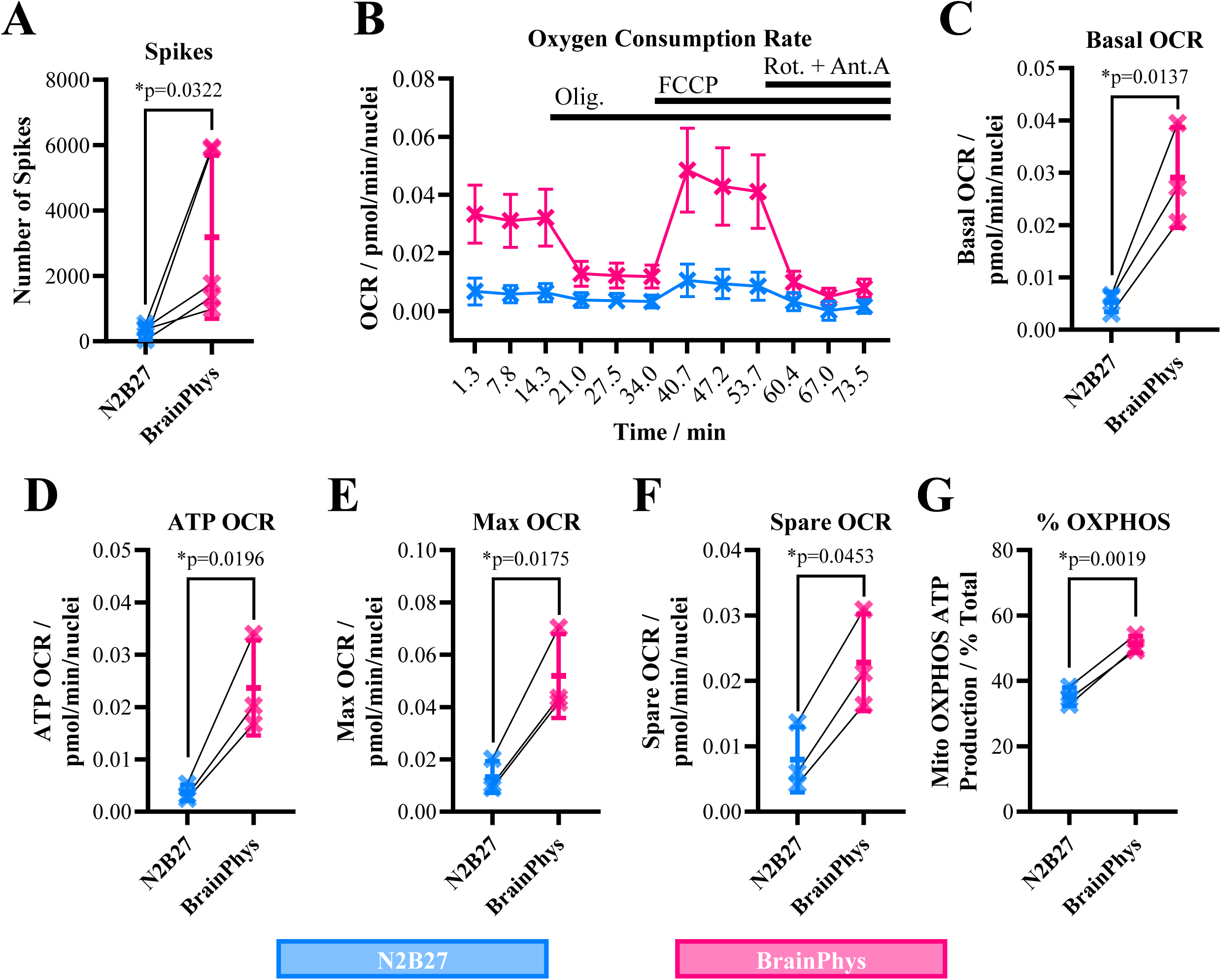
BrainPhys iNeurons are more electrically active and have more active mitochondrial oxidative phosphorylation. (**A**) Mean number of spikes detected per well (8x electrodes) of d24 N2B27 and BrainPhys iNeurons across a 15min MEA recording acquisition. (**B**) Representative OCR measurements made from d24 N2B27 and BrainPhys iNeurons following the sequential addition of 1µM oligomycin (Olig), 1µM FCCP, and 0.5µM equimolar mix of rotenone and antimycin-A (Rot.+Ant.A). Data are shown as the mean ±□SD from 15x technical replicate wells of a representative induction. (**C-F**) Measurements/calculations of basal (**C**), ATP (**D**), Maximal (**E**) and Spare (**F**) OCRs from (**B**). (**G**) % of ATP generated through mitochondrial OXPHOS calculated from OCR and ECAR measurements made following the sequential addition of 1.5µM oligomycin and 0.5µM equimolar rotenone+antimycin-A. With the exception of data presented in (**B**), data are shown as the mean with error bars□±□SD from biological replicates, with normalised values for each biological replicate shown as individual markers.

Moreover, under conditions that induce complete MMP collapse, such as 5µM O/A or 10µM CCCP (Extended Data Figure 4A-4B), reductions in PINK1 accumulation and pUb(Ser65) deposition are still observed in BrainPhys iNeurons (Extended Data Figure 5A-C). These data suggest that the reduction in PINK1 protein and PINK1-dependent pUb(Ser65) observed in BrainPhys iNeurons is not specific to mitochondrial stress induced PINK1 stabilisation but rather to an overall reduction in PINK1 protein availability.

**Figure 4.**
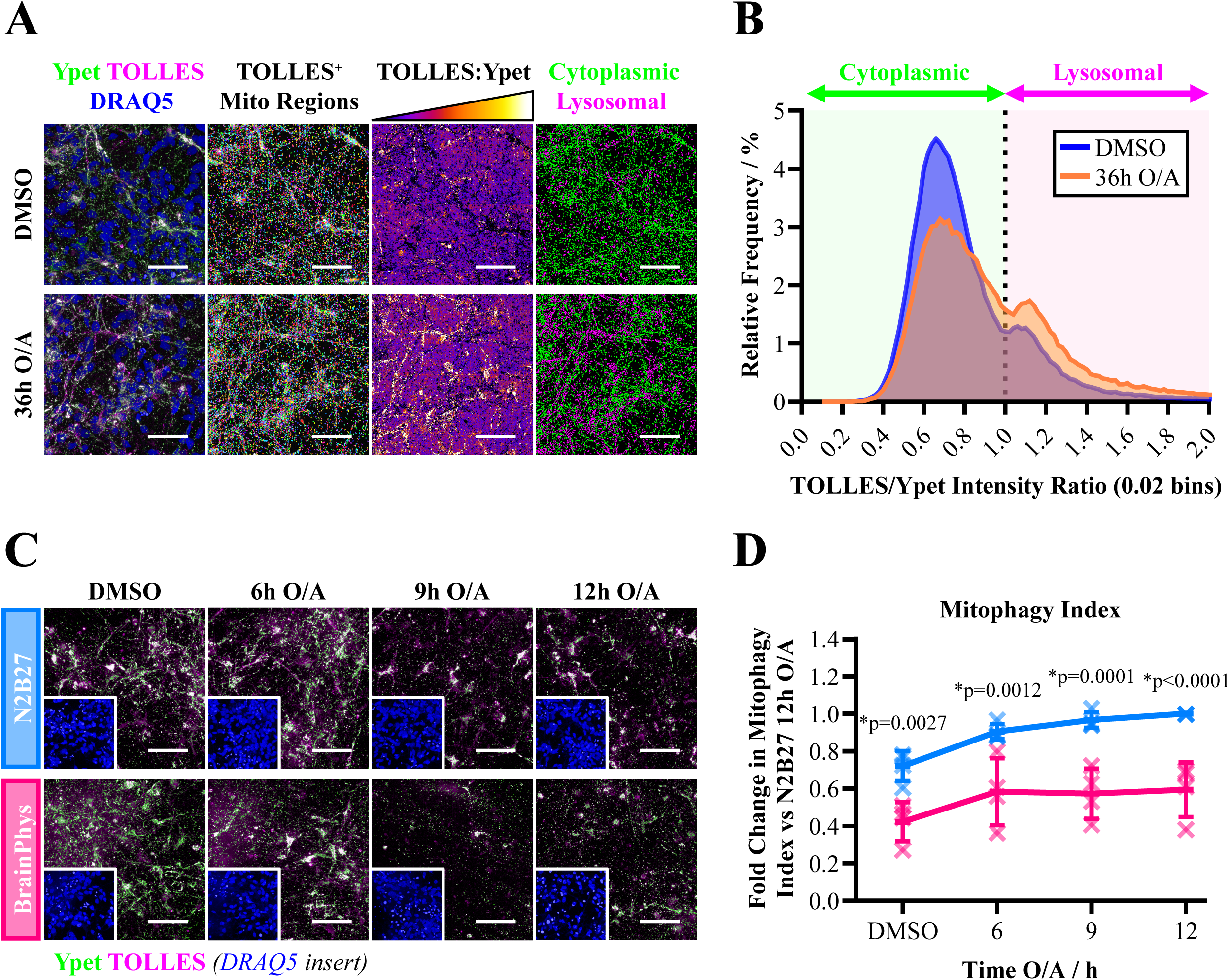
BrainPhys iNeurons show reduced basal and stress-induce mitophagy fluxes. **(A)** Overview of the mitoSRAI imaging analysis workflow and sensitive detection of lysosomal mitochondrial delivery in N2B27 iNeurons treated with 1 µM O/A for 36h. The first column shows fluorescence detection of TOLLES (magenta) and Ypet (green) alongside DRAQ5 counterstained nuclei (blue). The second column shows spot detection of individual TOLLES^+^ mitochondrial units. The third column shows a TOLLES:Ypet ratiometric image whereby increased signal is indicative of Ypet loss from TOLLES^+^ regions. The fourth column shows stratification of TOLLES^+^ mitochondrial units as cytoplasmic (low TOLLES:Ypet ratio / Ypet^+^) and lysosomal (high TOLLES:Ypet ratio / Ypet^-^). Scale bars = 50 µm. (**B**) Relative frequency histogram showing the TOLLES:Ypet ratio of individual TOLLES^+^ mitochondrial units as in (**A**). Data plotted are from a total of 30x 205µm x 205µm fields of view across 5x replicate wells of a single induction. (**C**) Representative confocal maximum intensity projections of TOLLES and Ypet fluorescence following treatment of d24 N2B27 and BrainPhys iNeurons with DMSO or 1 µM O/A. Insets show the DRAQ5 counterstained nuclei (blue) for the same field of view. Scale bars = 50 µm. (**D**) Quantification of the mitophagy index from (**C**) as the proportion of TOLLES^+^ mitochondrial area which is lysosomal (n=4 inductions, 6x 205 µm x 205µm fields of view per well, 5 experimental wells per condition, two-way ANOVA with Šidák post hoc correction). Data are shown as the mean with error bars□±□SD from biological replicates, with normalised values for each biological replicate shown as individual markers.

### Limitations in glucose are a major contributor to resistance of BrainPhys iNeurons initiating PINK1-dependent mitophagy

BrainPhys has a nearly 10x fold lower glucose concentration (2.5mM) [41] compared to N2B27 (21.25mM) which given previous data linking glucose-dependent metabolic fluxes to PINK1-dependent mitophagy [22,36,48,49] represents a potentially important underlying difference between N2B27 and BrainPhys in relation to the pUb(Ser65) phenotypes described here. To this end, a N2B27 media was formulated with a lower 2.5mM glucose concentration and higher 0.5mM pyruvate concentration, matching the respective concentrations of these metabolic substrates in BrainPhys. Conversely BrainPhys was supplemented with additional glucose to match that of N2B27. In line with glucose concentration contributing to the pUb(Ser65) phenotype, reducing the glucose concentration of N2B27 attenuated pUb(Ser65) deposition following 9h of O/A induced mitochondrial depolarisation (Figure 2F and 2G).

Conversely and even more strikingly, pUb(Ser65) was significantly elevated by supplementing BrainPhys iNeurons with higher glucose concentrations. While these data show that glucose limitation plays a major role in PINK1-mitophagy initiation, neither lowering the glucose in N2B27 nor increasing the glucose in BrainPhys fully explained the differences in pUb(Ser65) deposition observed between these media, with other differences in their constituents likely contributing. Assessments of FL-PINK1 following O/A-induced MMP depolarisation, and of Δ-PINK1 following MG132-dependent proteasomal inhibition, indicate that the glucose-dependency of pUb(Ser65) deposition is largely driven by the reduced PINK1 protein availability (Extended Data Figure 6A-6C).

### BrainPhys iNeurons are more bioenergetically active and have an increased reliance on mitochondrial OXPHOS

Given previous work linking reductions in PINK1-Parkin dependent mitophagy and the metabolic state of neurons in culture [22], the effect culturing iNeurons in BrainPhys has on neuronal electrical and metabolic activities was explored in more detail. Corroborating previous data [41], iNeurons cultured in BrainPhys showed an increase in basal electrical activity when compared to N2B27 iNeurons, as measured using the CytoView 96-well multi-electrode array (Figure 3A). In line with an increased bioenergetic demand associated with greater basal electrical activity, and a decreased concentration of glucose, basal oxygen consumption rates (OCR) were increased in BrainPhys iNeurons, as measured using the Seahorse extracellular flux analyser (Figure 3B-3C). Measuring OCR after adding oligomycin, FCCP and rotenone+antimycin-A (Figure 3B) allows the OCR contributing to ATP production via OXPHOS, maximal OCR and spare respiratory capacity to be calculated. In line with BrainPhys iNeurons showing an overall increase in mitochondrial respiration, measures of ATP production OCR, maximal OCR and spare respiratory capacity OCR were all increased when compared to N2B27 (Figure 3D-3F). These data suggest that BrainPhys neurons have an overall increase in mitochondrial OCR and respiration capacity, confirming observations made by others in murine cortical neurons cultured in BrainPhys vs neurobasal medium [50]. Using the Seahorse extracellular flux analyser to simultaneously measure extracellular acidification rate (ECAR) and OCR following the sequential addition of oligomycin and rotenone+antimycin-A, the proportion of cellular ATP generation through glycolysis and mitochondrial OXPHOS could be measured. In line with an increased reliance on mitochondrial OXPHOS, the proportion of ATP generated through mitochondrial OXPHOS rather than glycolysis was increased in BrainPhys iNeurons (Figure 3G). Together, these data suggest that glucose limitations of BrainPhys iNeurons are correlated with elevations in mitochondrial respiration capacity and a bioenergetic shift towards increased reliance on mitochondrial OXPHOS for neuronal ATP production. These measurements were performed in Seahorse XF DMEM (10mM glucose, 1mM pyruvate, 2mM L-glutamine). Therefore, the observed differences more likely reflect alterations in neuronal bioenergetic capacity rather than actual OCR/ECAR under N2B27 vs BrainPhys culture conditions.

### BrainPhys iNeurons show reductions in basal and stress induced mitophagy

Several probes have been created that allow mitochondria delivered to lysosomes (bona fide mitophagy) to be detected through changes in fluorescence. Among two of the most widely used are mitokeima [51] and mitoQC [52]. More recently a new tandem-fluorophore construct known as mitoSRAI has been generated which consists of a mitochondrial matrix targeted and lysosomal resistant cyan fluorophore known as Tolerance of Lysosomal Environments (TOLLES) joined to a lysosomal protease and acid sensitive yellow fluorophore known as Ypet [42]. The loss of Ypet fluorescence allows TOLLES^+^ mitochondrial units, which have been delivered to the lysosome (Ypet^-^) to be distinguished from those which remain cytoplasmic (Ypet^+^). mitoSRAI has several advantages over mitokeima and mitoQC including maintenance of lysosomal vs cytoplasmic signal upon all typical fixation strategies and lower contribution of proteasome-dependant changes in the fluorescence readout. For the first time, the mitoSRAI mitophagy probe assay was optimised for use in human neurons in order to understand whether the bioenergetic and PINK1-dependent pUb(Ser65) differences observed by culturing iNeurons in BrainPhys impact downstream lysosomal delivery of mitochondria.

Stable expression of mitoSRAI at the hPSC stage was first established through lentiviral transduction followed by iNeuron differentiation and culture as described above. Initial optimisations were performed in N2B27 iNeurons which robustly survived O/A-induced mitochondrial depolarisation for at least 36h (Figure 4A-4B). Confocal imaging of differentiated mitoSRAI iNeurons revealed that under basal conditions, the vast majority of TOLLES^+^ mitochondria also show Ypet fluorescence, indicating they are non-lysosomal. Following 36h treatment with O/A, a noticeable increase in the number of TOLLES^+^ mitochondria showing low levels of Ypet fluorescence were observed, in line with the delivery of depolarised mitochondria to the lysosomes and subsequent Ypet proteolysis. A number of different analysis strategies have been previously employed to quantify readouts from mitophagy probes in cultured neurons. These include sequentially defining the areas of cytoplasmic and lysosomal fluorescent signals, or measuring lysosomal and cytoplasmic fluorescence intensity ratios [28,29,38]. Due to the relatively rare nature of mitochondrial delivery to lysosomes in neuronal models including iNeurons, analyses have also been used to count the number of lysosomal mitochondrial units and/or number of neurons with lysosomal mitochondria [33,38]. Here a strategy to define the lysosomal vs total mitochondrial area was optimised, that would allow a mitophagy index which considers the entirety of the mitochondrial population to be calculated. Importantly, the TOLLES:Ypet intensity ratio metric used to define the lysosomal and cytoplasmic mitochondrial units gave a robust and reproducible identification of two populations, where a threshold to separate lysosomal (high TOLLES:Ypet ratio) and cytoplasmic (low TOLLES:Ypet ratio) mitochondrial units could be rationally placed (Figure 4B). A population of lysosomal mitochondria was detected in neuronal cultures even under basal conditions. Treatment with O/A for 36h increased the relative contribution of the lysosomal population, replacing the cytoplasmic population, as expected from the induction of PINK1-dependant mitophagy and subsequent delivery of mitochondria to lysosomes.

Using the stable mitoSRAI iNeuron system, the impact of BrainPhys culture conditions on basal and mitochondrial stress induced mitophagy was then explored. Further highlighting a greater dependence of BrainPhys iNeurons on mitochondrial function, O/A treatments lasting longer than 12h often led to complete neuronal death and/or mass detachment of cultures from the dish surface, limiting our ability to reproducibly obtain sufficient BrainPhys iNeuron material for mitophagy analyses. For this reason, 12h and shorter O/A treatment durations were utilised for assessing the impact of BrainPhys on iNeuron mitophagy.

Interestingly, BrainPhys iNeurons showed a smaller proportion of lysosomal mitochondria under control culture conditions (Figure 4C-4D), potentially indicative of an overall reduction in basal mitophagy. O/A-induced mitochondrial depolarisation resulted in an increase in the proportion of lysosomal mitochondria in both N2B27 and BrainPhys iNeuron cultures.

However, the increase in BrainPhys iNeurons was more modest and even after 12h of treatment, it had not reached the basal mitophagy levels observed in N2B27 iNeuron cultures (Figure 4C-4D). These data therefore suggest that BrainPhys iNeurons show overall reductions in mitophagy fluxes, both basally and following O/A-induced mitochondrial stress, with reductions in stress-induced mitophagy correlating with the attenuation in PINK1-dependent pUb(Ser65) deposition described above.

## DISCUSSION

Improvements in our understanding of PD genetics and biology of the mitophagy process have gone hand in hand since the initial discovery of causative loss of function mutations in the *PINK1* and *Parkin* genes in families with early-onset Mendelian PD. Since then, several other mitophagy-independent functions of PINK1 and Parkin have been described, including regulation of mitochondrial derived vesicle dynamics and organelle targeting [53,54], mitochondrial biogenesis [55], fission/fusion [56–59] and trafficking [60]. While it is clear PINK1 and Parkin are heavily implicated in mitochondrial quality control, the relevance of their involvement in mitophagy within human neurons has been met with some scepticism. Crucial to better understanding this will be the use of robust *in vitro* human neuron models.

Here, data highlights the importance of *in vitro* culture conditions, specifically medium composition, when exploring the role of PINK1 and Parkin in human neurons. Previous work has shown that the PINK1-Parkin mitophagy process is sensitive to cellular metabolic state, with cells favouring oxidative metabolism being resistant to PINK1-Parkin mitophagy [22,36,48,49]. Our results further support this observation, with BrainPhys iNeurons expressing endogenous levels of PINK1 and Parkin showing elevations in mitochondrial OXPHOS and reduced PINK1-Parkin dependant mitophagy initiation.

Previous work has correlated these metabolic relances for PINK1-dependent mitophagy on limitations in ATP availability following mitochondrial induced stress [49], with reversal of OXPHOS complex V appearing to be an important contributor to this phenotype in neurons [22]. Importantly, complex V reversal is unlikely to be contributing to the phenotypic differences described here independent of the media condition due to the co-application of oligomycin (Complex V inhibitor) with antimycin for depolarisation of the IMM.

A robust increase in the proportion of mitochondria present in lysosomal compartments was observed in N2B27 iNeuron cultures after O/A induced IMM depolarisation. However, this increase was notably more subtle in BrainPhys iNeurons. This aligns with the resistance of BrainPhys iNeurons for PINK1-dependent initiation of mitophagy under mitochondrial stress conditions. In addition, a population of lysosomal mitochondria was readily observed even under basal conditions, particularly in N2B27 iNeuron cultures. This corroborates previous findings that mitochondrial components are among the most prevalent contents of neuronal autophagosomes under basal conditions [61], and supports reports of basal mitophagy detection in iNeuron models cultured in Neurobasal base medium [33,38]. While lysosomal mitochondria were detected in BrainPhys iNeurons under basal conditions they were far less common, potentially highlighting an overall reduction in mitophagy fluxes generally.

Knockout mouse and *Drosophila* models suggest PINK1 and Parkin are not overly contributing to basal levels of mitophagy in most cells *in vivo*, including neurons [62–66]. This suggests that the differences in basal mitophagy of N2B27 and BrainPhys iNeurons are unlikely to be PINK1-dependent and instead through other PINK1-independent mitophagy mechanisms more predominant under healthy basal conditions [67].

This does not necessarily refute the functional importance of PINK1-Parkin dependent mitophagy to neurons however, which through their selectivity towards damaged and depolarised mitochondria will be likely playing a key role in mitochondrial homeostasis. This is particularly relevant when considering potential cumulative effects across the long duration of a human lifespan which is several magnitudes beyond typical *in vitro* culture durations such as that conducted here. Given overall reductions in basal turnover of mitochondria observed in BrainPhys iNeurons, it will be interesting to understand whether they are more vulnerable to accumulation of mitochondrial damage and associated oxidative stresses within such *in vitro* culture durations. Despite being more resistant to widespread PINK1-dependant mitophagy initiation, it could be speculated that the PINK1-Parkin mitophagy process is even more important under such conditions.

These data leave open the question as to what media choices and culture conditions should be employed when probing human neuron mitochondrial homeostasis. The better recapitulation of physiological conditions with BrainPhys would certainly be a major argument for its use. Detailed mechanistic studies using this culture base will be important for furthering our understanding of human neuron biology, including the underlying importance, mechanisms and regulation of mitophagy pathways. In a similar way that researchers continue to utilise Parkin overexpression for mechanistic insights into PINK1-Parkin biology, the increased activities of both PINK1-dependent and basal mitophagy processes in N2B27 based medium does however lend support to its continued use. It is anticipated that in specific situations discoveries might more readily be made in N2B27 cultures which can then be validated in more physiological BrainPhys cultures.

In all, these data highlight the importance of acknowledging differences which exist in *in vitro* models when exploring and interpreting the importance of mitophagy in human neurons, and provide further insight into the PINK1-Parkin dependent mitophagy process in human neurons.

## METHODS

The following detailed protocols can be found at protocols.io:

hPSC Culture, iNeuron Differentiation and Culture in N2B27 vs BrainPhys for Immunofluorescence and Biochemistry Assessments of Mitophagy (https://dx.doi.org/10.17504/protocols.io.5qpvo36xxv4o/v3);

Immunofluorescence Staining for Assessing pUb(Ser65) Accumulation in iNeurons (https://dx.doi.org/10.17504/protocols.io.eq2ly6okpgx9/v1);

Immunoblot Assessments of PINK1-Dependent Mitophagy in iNeurons (https://dx.doi.org/10.17504/protocols.io.6qpvr9bj2vmk/v1);

iNeuron RNA-Extraction and RT-qPCR (https://dx.doi.org/10.17504/protocols.io.ewov1drryvr2/v1);

Measuring Mitochondrial Membrane Potential in iNeurons Through Live-Cell TMRM Imaging (https://dx.doi.org/10.17504/protocols.io.14egn1yjqv5d/v1);

Seahorse OCR/ECAR, MitoStress Test and ATP-Rate Test for iNeurons (https://dx.doi.org/10.17504/protocols.io.5qpvo36xxv4o/v1);

Measurement of iNeuron electrical activity using Axion Biosystems Maestro Pro MEA (https://dx.doi.org/10.17504/protocols.io.5qpvo36xxv4o/v2);

Detection of Lysosomal Delivery of Mitochondria in iNeurons Using MitoSRAI Reporter (https://dx.doi.org/10.17504/protocols.io.261ged7qjv47/v1).

hPSC Culture and iNeuron Differentiation:

A WTC11 hPSC line harbouring a TET-ON system at the AAVS1 safe-harbour locus for overexpression of murine Ngn2 was a kind gift from the laboratories of M.E. Ward and M. Kampmann, and have been published elsewhere [43,44]. hPSCs were cultured in mTeSR Plus medium (StemCell Technologies) on Geltrex (Thermo Fisher) coated culture dishes and maintained in a humidified incubator at 37°C and 5%/95% CO_2_/air mixture.

hPSCs were differentiated into iNeurons by adapting previously published protocols [35,43]. Cells were maintained in a humidified incubator at 37°C and 5%/95% CO_2_/air mixture throughout. On d-1 hPSCs were dissociated into a single cell-suspension using TrypLE Express (Gibco) and seeded onto geltrex coated 6-well culture dishes in mTeSR Plus medium supplemented with 10 µM Y27632 Rho-kinase inhibitor (ROCKi; MedChem, HY-10071) at a seeding density of 4.5×10^5^ cells per well. The next day (d0) medium was changed for induction medium consisting of DMEM/F12-HEPES supplemented with 1x N2 supplement and 1x non-essential amino acids (NEAA) (all Gibco) and 2µg/ml doxycycline (Sigma). A full medium change was performed 24h (d1) and 48h (d2) later with fresh induction medium. On d3 iNeuron neural progenitor cells (NPCs) were dissociated into a single cell suspension using accutase (Sigma) and seeded in N2B27 medium or BrainPhys medium. The concentrations of glucose, pyruvate and amino acids within the media formulations used throughout this study are summarised in Table 1. N2B27 medium consisted of a 1:1 mix of DMEM/F12-HEPES and neurobasal supplemented with 0.5x N2 supplement, 0.5x B27 Supplement, 0.5x NEAA, 0.5x Glutamax, 45 µM β-mercaptoethanol (all Gibco) and 2.5µg/ml insulin (Sigma). BrainPhys medium consisted of the BrainPhys base (Stem Cell Technologies), supplemented with 1x B27 Supplement, 10ng/ml Brain-derived neurotrophic factor (BDNF; Peprotech) and 10ng/ml Neurotrophin-3 (NT-3; Peprotech). Details on seeding densities are described in the specific sections below. For experiments controlling the glucose and pyruvate concentrations of N2B27, a 1:1 mix of glucose, glutamine and pyruvate free DMEM (Gibco, Cat# A1443001) and Neurobasal-A (Gibco, Cat# A2477501) were supplemented with 2mM Glutamax, 0.5x N2 supplement, 0.5x B27 Supplement, 0.5x NEAA, 45 µM β-mercaptoethanol (all Gibco) and 2.5µg/ml insulin (Sigma). This base was supplemented with 21.25mM glucose (typical high concentration) and 0.36mM pyruvate or alternatively lower 2.5mM glucose and 0.5mM puryvate (all Gibco). High glucose BrainPhys was also prepared by supplementing the complete BrainPhys medium described above to a final glucose concentration of 21.25mM. Half media changes was performed 3 days after seeding (d6) and every 3-4 days thereafter. A half media change was performed on d23, 24h prior to collection for experimental assays. Each biological replicate represents an independent iNeuron differentiation from a separate induction batch. For each biological replicate experimental comparisons will be made between samples differentiated alongside one another and experimentally assayed simultaneously.

### Dual SMADi Cortical Neuron Differentiation

Cortical neurons were differentiated from 3 different control hPSC lines/genotypes (RBi001-A, RRID:CVCL_9S35; SIGi001-A-1, RRID:CVCL_EF83; WTSIi019-B, RRID:CVCL_AE22) through well-established dual SMAD inhibition strategies as described previously[68–70]. Briefly, hPSCs were grown to 100% confluency before performing a full media change to neural induction media consisting of N2B27 (as described above) additionally supplemented with 10□μM SB431542 (Tocris) and 1□μM dorsomorphin (Tocris) (d0). Full neural induction media changes were performed daily until d10-12, at which point the neuroepithelial layer was detached with 10mg/ml dispase (Invitrogen). The neuroepithelial layer was dissociated into clumps of cells which were then replated in N2B27 onto geltrex or laminin (Sigma) coated dishes, with N2B27 media changes performed every 3-4 days. On d35 NPCs were dissociated into a single cell suspension using accutase, and seeded into 24-well (1×10^5^ cells per well) or 12-well (5×10^5^ cells per well) in N2B27 media. Half media changes with N2B27 were performed every 3-4 days until was changed daily until d55 at which point a full media change was performed with N2B27 or BrainPhys. Cells were then cultured for additional 3 weeks with N2B27 vs BrainPhys half media changes every 3-4 days. A half media change was performed on d75 1h prior to beginning a 24h O/A time-course which was collected on d76.

### Whole-cell lysis and Immunoblotting

For whole-cell protein lysates, 6×10^5^ d3 iNeuron NPCs were seeded into each well of a geltrex coated 12-well plate and cultured until d24 as described above. Cells were treated with 1 µM equimolar combination of oligomycin and antimycin (O/A) (Sigma), 0.5 µM O/A, 5 µM O/A, 1 µM rotenone (Sigma), 5 µM rotenone, 10 µM CCCP (Sigma), 10 µM CCCP + 1 µM Oligomycin (Sigma) and/or 10 µM MG132 (MedchemExpress) prior to collection in ice-cold RIPA-lysis buffer consisting of: 50mM Tris-HCl (pH=7.4), 150mM NaCl, 1% w/v triton-X-100, 0.5% sodium deoxycholate, 0.1% w/v sodium dodecyl sulphate (SDS), 1x PhosSTOP phosphatase inhibitors and 1x cOmplete™, Mini, EDTA-free Protease Inhibitors (all Sigma). Equal amounts of protein lysate (typically >10µg) were separated through SDS polyacrylamide gel electrophoresis (PAGE) using 4-12% NuPAGE Bis-Tris Mini Protein Gels and XCell SureLock Mini Cell (Thermo Fisher Scientific), before transfer to 0.45 µm Immobilon®-FL PVDF membrane (Sigma) using Mini Trans-Blot Cell (Bio-Rad). Blots were blocked with 5% w/v milk in PBS with 0.01% tween-20 (PBST; 1h, room temperature (RT); Sigma). Blots were incubated with the following primary antibodies diluted in 3% w/v milk in PBS-T (4°C, overnight): rabbit anti-pUb(Ser65) IgG (1:1000; Cell Signaling Technology Cat# 62802, RRID:AB_2799632), rabbit anti-PINK1 (1:1000; Takeda, RRID:AB_3661827), rabbit anti-Parkin (1:500; Invitrogen Cat# 702785, RRID:AB_2942251), mouse anti-TIM23 IgG2a (1:1000; BD Biosciences Cat# 611223, RRID:AB_398755), mouse anti-GAPDH IgG2b (1:10000; Abcam Cat# ab110305, RRID:AB_10861081) and mouse anti-β3-Tubulin Mouse IgG2a (1:10000; (BioLegend Cat# 801201, RRID:AB_2313773). The generation of rabbit anti-PINK1 IgG antibody (RRID:AB_3661827) has been described previously[71] and is available on reasonable request to the corresponding authors. Blots were then washed 3x with PBST prior to incubation with fluorescently conjugated secondary antibodies diluted in PBST supplemented with 0.02% SDS (2h, RT): donkey anti-mouse IgG (H+L) IRDye 680LT (1:20000; LI-COR Biosciences Cat# 926-68022, RRID:AB_10715072) and donkey anti-rabbit IgG (H+L) IRDye 800CW (1:20000 LI-COR Biosciences Cat# 926-32213, RRID:AB_621848). Blots were washed 3x with PBST and 1x with PBS prior to near-infrared (NIR) detection using an Odyssey CLx imager (LI-COR Biosciences, RRID:SCR_014579). NIR signal integrated densities were quantified using Image Studio Lite v5.2 analysis software (LI-COR Biosciences, RRID: SCR_013715). GAPDH integrated density acted as a loading normalisation control.

### Immunofluorescence and Confocal Imaging

For immunofluorescence confocal imaging experiments, 5×10^4^ d3 iNeuron NPCs were seeded into each well of a geltrex coated 96-well PhenoPlate (Revvity) and cultured until d24 as described above. Cells were treated with 1 µM O/A and then fixed with 4% formaldehyde (Sigma; 20min, RT). Cells were washed 3x with phosphate buffered saline (PBS; Fisher Scientific) prior to blocking and permeabilisation with 1x PBS supplemented with 10% v/v foetal bovine serum (FBS) and 0.25% w/v triton-x-100 (both Gibco; 1h, RT). Cells were then incubated with the following primary antibodies diluted in blocking buffer (10% v/v FBS in 1x PBS, 3h, RT): rabbit anti-pUb(Ser65) IgG (1:1000; Cell Signaling Technology Cat# 62802, RRID:AB_2799632), mouse anti-TOM20 IgG2a (1:1000; Santa Cruz Biotechnology Cat# sc-17764, RRID:AB_628381), chicken anti-MAP2 IgY (1:5000; Millipore Cat# AB5543, RRID:AB_571049). Cells were then washed 3x with PBS prior to incubation with fluorescently conjugated secondary antibodies diluted in blocking buffer (2h, RT): goat anti-rabbit IgG (H+L) Alexa Fluor 488 (1:2000; Thermo Fisher Scientific Cat# A-11008, RRID:AB_143165), goat anti-mouse IgG (H+L) Alexa Fluor 568 (1:2000; Thermo Fisher Scientific Cat# A-11004 (H+L), RRID:AB_2534072), goat anti-chicken IgY (H+L) Alexa Fluor 647 (1:2000, Thermo Fisher Scientific Cat# A-21449, RRID:AB_2535866). 10 µM Hoechst 33342 (Thermo Fisher Scientific Cat# 62249) was included in the secondary antibody dilution to counterstain nuclei. Fluorescence of each channel was captured sequentially with non-saturating dwell times and the following excitation lasers and emission filters: Hoechst 33342 (Ex/Em: 375nm/435-515nm), Alexa Fluor 488 (Ex/Em: 488nm/500-550nm), Alexa Fluor 568 (Ex/Em: 561nm/570-630nm), Alexa Fluor 647 (Ex/Em: 640nm/650-760nm). For each experimental well 1 µm thick z-slices encompassing the entirety of the sample depth were captured every 1 µm across 6x 323×323 µm fields of view. The mean pixel intensity of pUb(Ser65) immunofluorescence (Alexa Fluor 488) within the MAP2^+^ area mask (Alexa Fluor 647) was calculated from maximum intensity z-projections using Columbus image analysis software (Revvity). Each biological data point represents the mean of 4-6 experimental wells for each condition.

### RNA Extraction and RT-qPCR

For RNA, 6×10^5^ d3 iNeuron NPCs were seeded into each well of a geltrex coated 12-well plate and cultured until d24 as described above. Total RNA was extracted using the Monarch Total RNA Miniprep Kit (New England Bioscience) with inclusion of the optional on-column DNAse treatment and quantified using a NanoDrop One Spectrophotometer (Thermo Fisher Scientific). 100 ng of RNA was reverse transcribed in a 10 µl reaction with 2.5 U/µl SuperScript IV reverse transcriptase, 2.5 µM random hexamers, 0.5 mM dNTPs, 5 mM DTT and 2 U/µl RNAseOUT (all Invitrogen). The cDNA product was then diluted such that 100 ng of reverse transcribed RNA would be in a 120µl final volume (i.e. 0.83 ng/µl). 3.33 ng cDNA was then subjected to real-time quantitative PCR (qPCR) using 1× Power SYBR Green Master Mix (Applied Biosystems) and 500 nM gene specific primer pairs on a QuantStudio™ 7 Flex Real-Time PCR System (Applied Biosystems, RRID:SCR_020245). At least 2× technical replicates were performed for each sample and gene target combination, and an RT^-^ control for all samples and gene target combinations was performed alongside. Relative *PINK1* mRNA expression levels were calculated using the 2^−ΔΔCt^ method and *UBC* as the house-keeping gene.

### TMRM Measurements of iNeuron Mitochondrial Membrane Potential

For confocal imaging of TMRM intensity, 5×10^4^ d3 mitoSRAI iNeuron NPCs were seeded into each well of a geltrex coated 96-well PhenoPlate (Revvity) and cultured until d24 as described above. Cells were treated with 0.5 µM O/A, 1 µM O/A, 5 µM O/A, 1 µM rotenone, 5 µM rotenone, 10 µM CCCP or 10 µM CCCP + 1 µM Oligomycin for 1h, followed by addition of 25 nM TMRM (Sigma) for an additional 1h. Confocal images were captured using the Opera Phenix High-Content Screening System and 63x water objective, with TMRM remaining in the media throughout imaging acquisition. Fluorescence of each channel was captured sequentially with non-saturating dwell times and the following excitation lasers and emission filters: Ypet (Ex/Em: 488nm/500-550nm) and TMRM (Ex/Em: Ex/Em: 561nm/570-630nm). For each experimental well 1µm thick z-slices encompassing the entirety of the sample depth were captured every 1µm across 5x 205µm x 205µm fields of view. Columbus image analysis software was used to calculate the mean TMRM intensity within YPet^+^ mitochondrial regions. Each biological data point represents the mean of 3 experimental wells for each condition.

### MEA Measurements of iNeuron Electrical Activity

The electrode array of CytoView MEA 96-well plates was spot coated with 2x concentrated geltrex by adding a 8ul droplet to the centre of each well. 5×10^4^ d3 iNeuron NPCs were then seeded onto the geltrex coated electrodes in an 8ul droplet of N2B27. 2-3h later the wells were then flooded with 150µl N2B27 or BrainPhys and cultured until d24 as described above. iNeuron electrophysiological activity was recorded using an Mastero Pro MEA machine (Axion Biosystems) at 12.5kHz sampling frequency and spikes corresponding to individual neuronal action potentials detected and quantified using AxIS Navigator software (Axion Biosystems). Each biological data point represents the mean number of spikes detected within a 15min recording across all 8 electrodes for each of 10 experimental wells per condition.

### Measurements of iNeuron Mitochondrial Respiration and Metabolic Fluxes

2×10^4^ d3 iNeuron NPCs were seeded into each well of a geltrex coated Seahorse XF96 plate and cultured until d24 as described above. iNeurons were preincubated in Seahorse XF DMEM supplemented with 10mM Glucose, 1mM pyruvate and 2mM L-glutamine for 1h, prior to a full fresh change just before conducting the first seahorse measurements. Oxygen Consumption Rate (OCR) and Extracellular Acidification Rate (ECAR) were recorded for 3min every 6.5min using the Seahorse XFE96 Bioanalyser (Agilent). Sequential injection of mitochondrial toxins oligomycin, FCCP and rotenone+antimycin (Agilent and Sigma) permit measurements of ATP production via OXPHOS, maximal OCR and calculations associated with spare respiratory capacity to be made using Wave v2.6.3 analysis software as part of the mito stress test (Agilent). Sequential injection of mitochondrial toxins oligomycin and rotenone+antimycin-A permit calculations of the proportion of cellular ATP generation through glycolysis and mitochondrial OXPHOS to be made using Wave v2.6.3 analysis software as part of the XF Real-Time ATP Rate Assay (Agilent). Following OCR/ECAR measurements, nuclei were counterstained with 10 µM Hoechst 33342 and each well tile-imaged using the Opera Phenix High-Content Screening System (Perkin Elmer) and 20x water objective (Ex/Em: 375nm/435-515nm). Total number of nuclei for each well were calculated using Columbus image analysis software and used to normalise all OCR and ECAR measurements in the Wave v2.6.3 analysis software. Each biological data point represents the mean of 15 experimental wells for each condition.

### MitoSRAI Measurements of iNeuron Mitophagy

Generation of the pLVX-EF1α-mitoSRAI-IRES-Puro plasmid and second-generation lentiviral particles have been previously described [72]. Stable expression of mitoSRAI was established through reverse transduction of 1×10^6^ iNeuron hPSCs in mTeSR Plus medium supplemented with 5 µg/ml polybrene (Sigma) and 10 µM ROCKi. Stably expressing cells were selected with 1 µg/ml puromycin (M P BIOMEDICALS UK) for >3 weeks prior to any iNeuron inductions. Puromycin was maintained during routine culture but withdrawn when seeding for iNeuron differentiation. hPSCs were cultured and differentiated as described above for immunofluorescence. iNeurons were cultured until d24 as described above, treated with 1µM O/A and then fixed with 4% formaldehyde (20min, RT). Cells were then washed 3x with PBS before counterstaining of nuclei with 2.5µM DRAQ5 diluted in PBS (2h, RT). Cells were washed 3x with PBS before confocal image acquisition using the Opera Phenix High-Content Screening System and 63x water objective. Fluorescence of each channel was captured sequentially with non-saturating dwell times and the following excitation lasers and emission filters: TOLLES (Ex/Em: 425nm/435-515nm), Ypet (Ex/Em: 488nm/500-550nm), and DRAQ5 (Ex/Em: 640nm/650-760nm). For each experimental well 1µm thick z-slices encompassing the entirety of the sample depth were captured every 1µm across 6x 205µm x 205µm fields of view. Columbus image analysis software was used to generate a mitophagy index from maximum intensity z-projections as follows. Mitochondrial units were first defined using the spot detection tool and TOLLES fluorescence signal. Mitochondrial units were then stratified into cytoplasmic and lysosomal based on the TOLLES:Ypet intensity ratio (High TOLLES:Ypet ratio classed as Ypet^-^ and lysosomal). The mitophagy index for each well was calculated as: 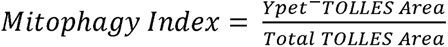. Each biological data point represents the mean of 5 experimental wells for each condition.

### Graphing and Statistical Analysis

GraphPad Prism 10 (RRID:SCR_002798) was used for generation of graphs and statistical analyses. Data were subjected to two-way ANOVA with Šidák *post hoc* correction for multiple comparisons or unpaired two-tailed t-test. Data are shown as the mean with error bars□±□standard deviation (SD) from biological replicates (independent neuronal inductions), with normalised values for each biological replicate shown as individual markers.

### Availability Statement

The datasets, software, protocols, and lab materials used and/or generated in this study are listed in a Key Resource Table alongside their persistent identifiers at https://dx.doi.org/10.5522/04/29064815. All datasets are made available on reasonable request or already deposited and available at https://dx.doi.org/10.5522/04/29064815. No code was generated for this study; all data cleaning, preprocessing, analysis, and visualization was performed using Prism 10:0:0 GraphPad (RRID:SCR_002798), LI-COR Image Studio Software (RRID:SCR_015795), Harmony v4.8 Software (RRID:SCR_018809), Columbus Image Data Storage and Analysis System (PerkinElmer https://www.perkinelmer.com/product/image-data-storage-and-analysis-system-columbus), Axion Integrated Studio (AxIS) Navigator (RRID:SCR_016308) and Seahorse Wave Software (RRID:SCR_024491). This manuscript was first deposited as a preprint which can be found at https://dx.doi.org/10.1101/2025.04.01.646648 [73].

## Supporting information

Key Resource Table

Tabular Data

## ACKNOWLEDGEMENTS

This research was funded in whole or in part by Aligning Science Across Parkinson’s (grant number ASAP 000478) through the Michael J. Fox Foundation for Parkinson’s Research (MJFF). For the purpose of open access, the author has applied a CC BY public copyright licence to all Author Accepted Manuscripts arising from this submission. B.O’C. is supported by a Guarantors of Brain Postdoctoral Non-Clinical Fellowship. D.M. was supported by an MRC CASE studentship (MR/P016677/1). K.C. and D.S. were supported by Eisai, working within the Eisai:UCL Therapeutic Innovation Group (TIG) with funding to H.P-F. M.S. was funded by MRC MR/N026004/1. N.B. is supported by The Motor Neuron Disease Association fellowship: Birsa/Oct21/976-799. S.W. and C.A. are supported by the NIHR UCL Hospitals Biomedical Research Centre.

## ABBREVIATIONS

DMEM: Dulbecco’s Modified Eagle Medium
hPSCs: human pluripotent stem cells
IMM: inner mitochondrial membrane
iNeurons: induced neurons
MMP: mitochondrial membrane potential
O/A: oligomycin-antimycin
OMM: outer mitochondrial membrane
PARL: presenilin-associated rhomboid-like protease
PD: Parkinson’s disease
PINK1: PTEN-induced kinase 1
TOM: translocase of the outer membrane.

---

**Extended Data Figure 1 – The intensity of pUb(Ser65) within MAP2^+^ iNeurons is reduced when cultured in BrainPhys.** (**A**) Representative confocal maximum intensity projections of pUb(Ser65) immunofluorescence (green) following treatment of d24 N2B27 and BrainPhys iNeurons with DMSO or 1 µM O/A. Insets show the Hoechst 33342 counterstained nuclei (blue) and MAP2 immunofluorescence (magenta) for the same field of view. Scale bars = 50 µm. (**F**) Quantification of mean pUb(Ser65) intensity within MAP2^+^ regions in (**E**) (n=5 inductions, 6x 323 µm x 323 µm fields of view per well, 4-6 experimental wells per condition, two-way ANOVA with Šidák *post hoc* correction).

Data are shown as the mean with error bars□± SD from biological replicates, with normalised values for each biological replicate shown as individual markers.

**Extended Data Figure 2 – Cortical Neurons differentiated through dual SMADi show impairments in PINK1-dependent mitophagy initiation when cultured in BrainPhys.** (**A**) Representative immunoblots of d76 cortical neurons cultured in N2B27 vs BrainPhys medium for 3 weeks prior to collection, and treated with 1 µM O/A over a 24h time-course. (**B**) Quantification of pUb(Ser65) from (**A**) (n=3 inductions from each from a separate control hPSC line/genotype, two-way ANOVA with Šidák *post hoc* correction).

Data are shown as the mean with error bars□± SD from biological replicates, with normalised values for each biological replicate shown as individual markers.

**Extended Data Figure 3 – Reductions in O/A induced FL-PINK1 protein availability in BrainPhys iNeurons do not appear to be linked with enhanced cleavage of PINK1 (**Δ**PINK1) and its subsequent degradation.** (**A**) Representative immunoblots of d24 iNeurons cultured in N2B27 vs BrainPhys medium and treated with 1 µM O/A, 10 µM MG132 or co-application of O/A + MG132 for 9h. (**B-D**) Quantification of full-length PINK1 (FL-PINK1) (**B**), PARL-cleaved PINK1 (ΔPINK1) (**C**) and calculation of the FL-PINK1:ΔPINK1 ratio from (**A**) (n=4 inductions, two-way ANOVA with Šidák *post hoc* correction).

Data are shown as the mean with error bars□±□SD from biological replicates, with normalised values for each biological replicate shown as individual markers.

**Extended Data Figure 4 – BrainPhys iNeurons have a more depolarised basal mitochondrial membrane potential and show comparable levels of mitochondrial membrane potential following application of a range of different mitochondrial toxins and doses.** (**A**) Representative confocal maximum intensity projections of d24 N2B27 and BrainPhys iNeurons stained with 25 nM TMRM, and pretreated for 2h with DMSO or various mitochondrial toxins (number indicates [toxin] in µM; ROT = rotenone, O = Oligomycin). Insets show the YPet^+^ mitochondria used for defining TMRM measurement areas for the same field of view. Scale bars = 50 µm. (**B**) Quantification of mean TMRM intensity within YPet^+^ mitochondrial regions in (**A**) (n=3 inductions, 5x 205 µm x 205 µm fields of view per well, 3 experimental wells per condition, two-way ANOVA with Šidák *post hoc* correction).

Data are shown as the mean with error bars□± SD from biological replicates, with normalised values for each biological replicate shown as individual markers.

**Extended Data Figure 5 – BrainPhys iNeurons show impairments in pUb(Ser65) deposition and PINK1 accumulation following application of a range of different mitochondrial toxins and doses.** (**A**) Representative immunoblots of d24 iNeurons cultured in N2B27 vs BrainPhys medium and treated for 9h with DMSO or various mitochondrial toxins (number indicates [toxin] in mM; ROT = rotenone, O = Oligomycin). (**B-C**) Quantification of pUb(Ser65) (**B**) and PINK1 (**C**) from (**A**) (n=3 inductions, two-way ANOVA with Šidák *post hoc* correction).

Data are shown as the mean with error bars□± SD from biological replicates, with normalised values for each biological replicate shown as individual markers.

**Extended Data Figure 6 – Differences in PINK1 protein availability between N2B27 and BrainPhys iNeurons is at least partly due to differences in glucose availability.** (**A**) Representative immunoblots of d24 iNeurons cultured in N2B27 vs BrainPhys medium with low 2.5mM glucose or high 21.25mM glucose and treated with 1 µM O/A or 10 µM MG132 for 9h. (**B-C**) Quantification of full-length PINK1 (FL-PINK1) (**B**) and PARL-cleaved PINK1 (ΔPINK1) (**C**) from (**A**) (n=4 inductions, two-way ANOVA with Šidák *post hoc* correction).

Data are shown as the mean with error bars□±□SD from biological replicates, with normalised values for each biological replicate shown as individual markers.

