## Supplementary figures and images for "Glucose concentration of neuronal media formulations influences PINK1-dependent mitophagy in human iNeurons"

### Extended Data Figure 1

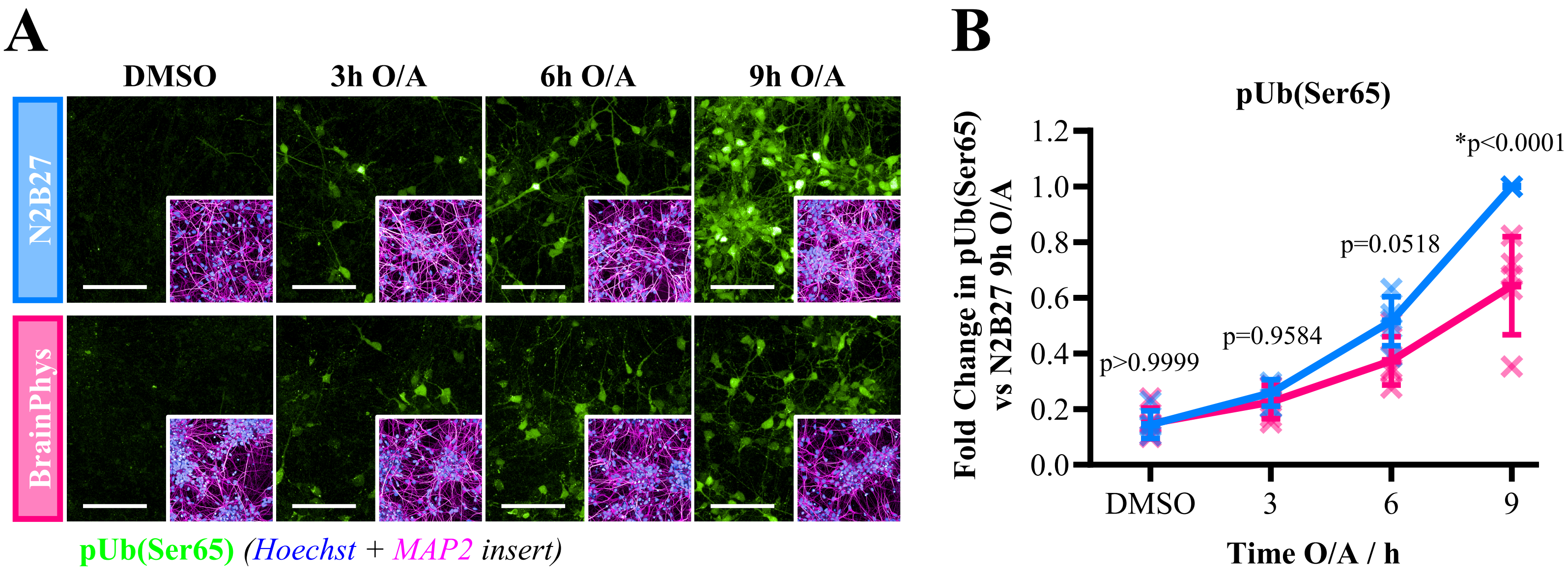

### Extended Data Figure 2

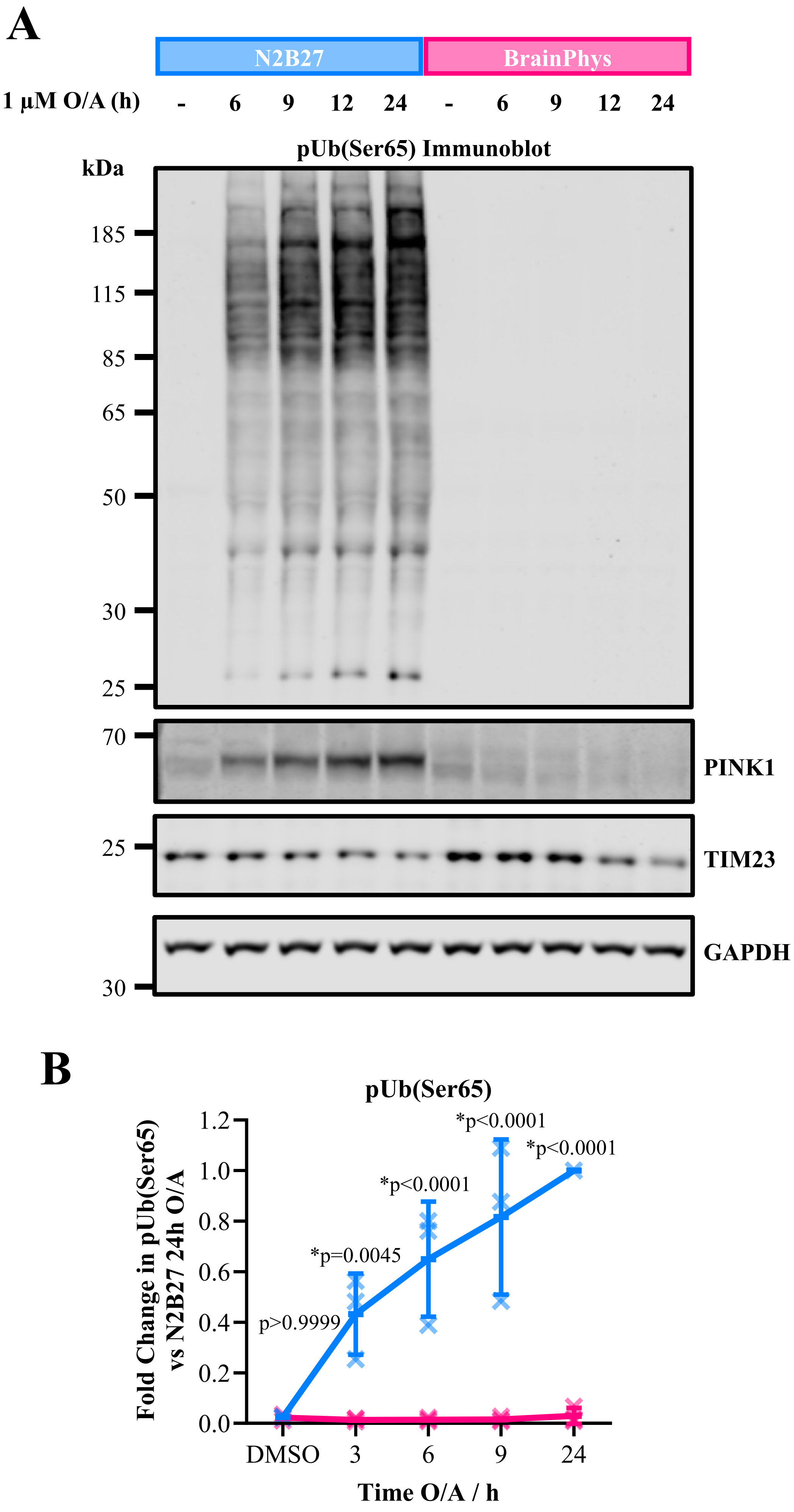

### Extended Data Figure 3

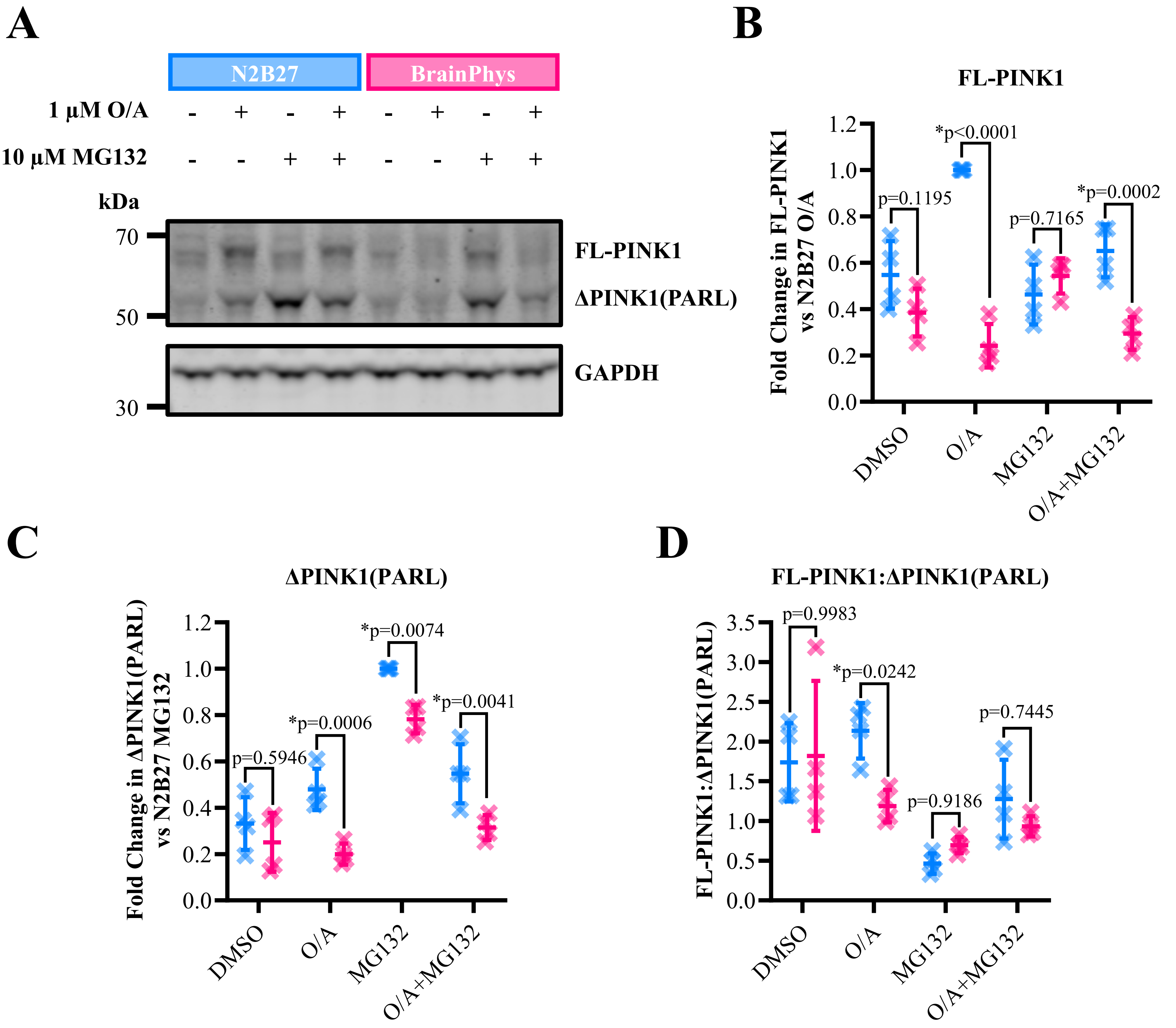

### Extended Data Figure 4

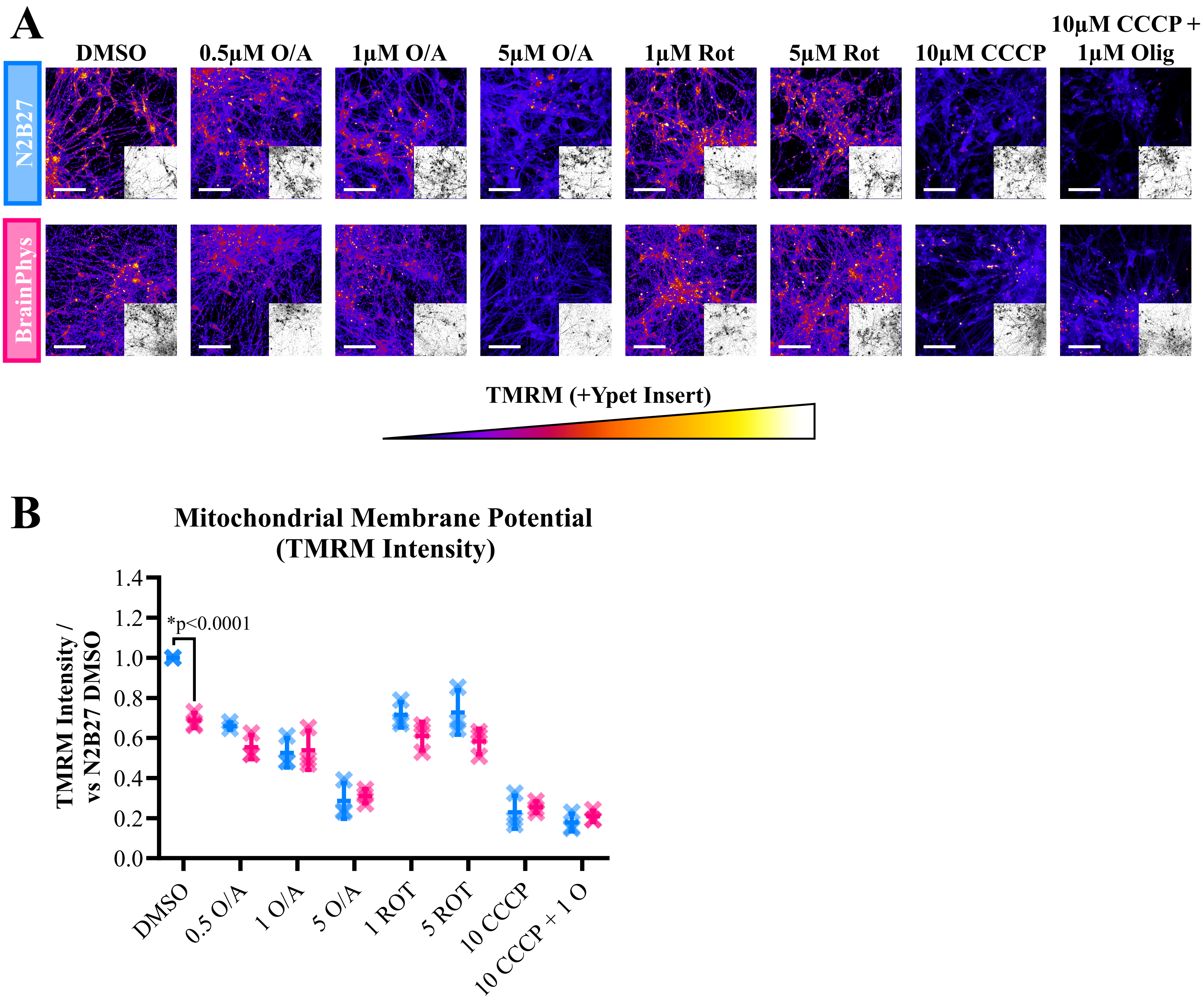

### Extended Data Figure 5

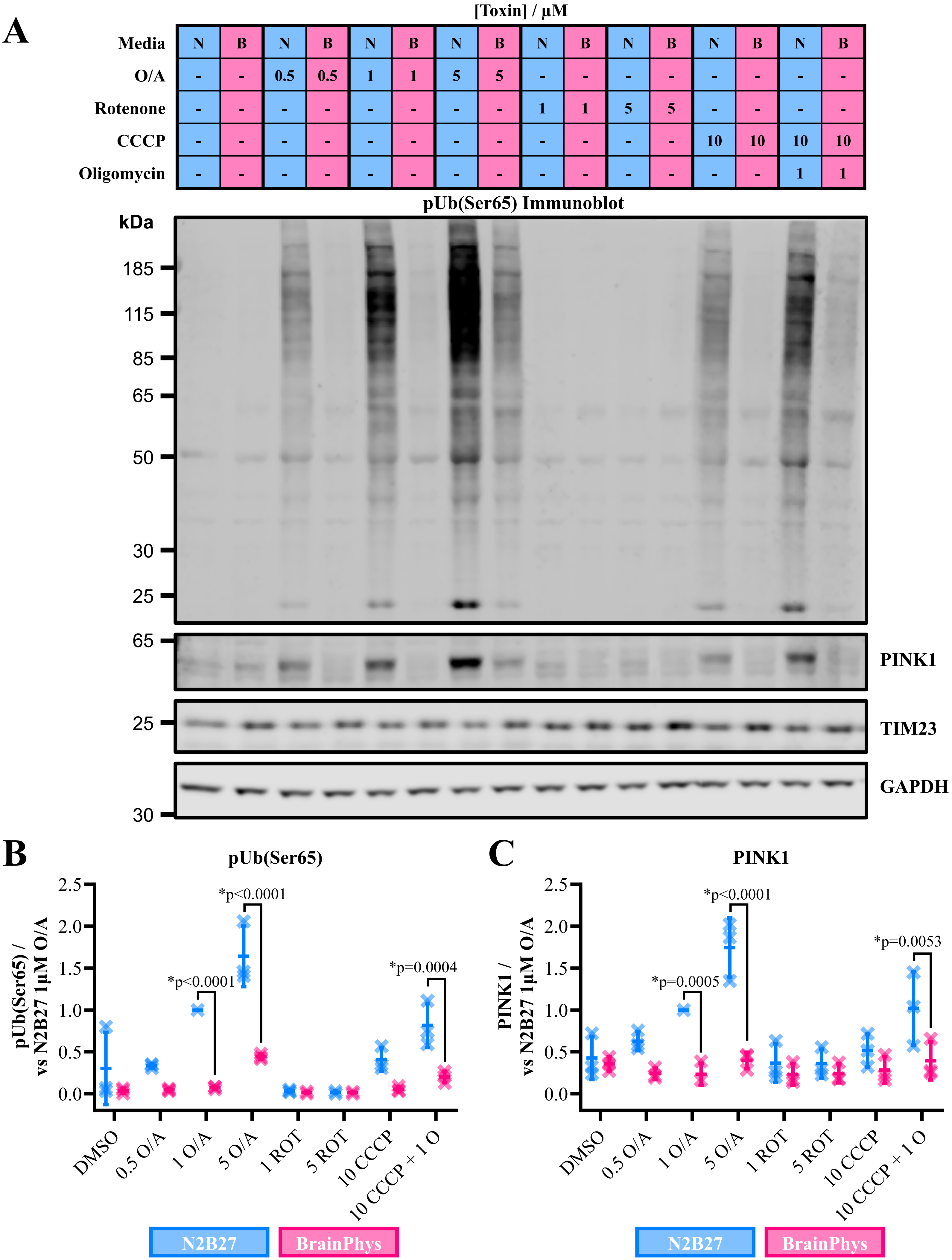

### Extended Data Figure 6

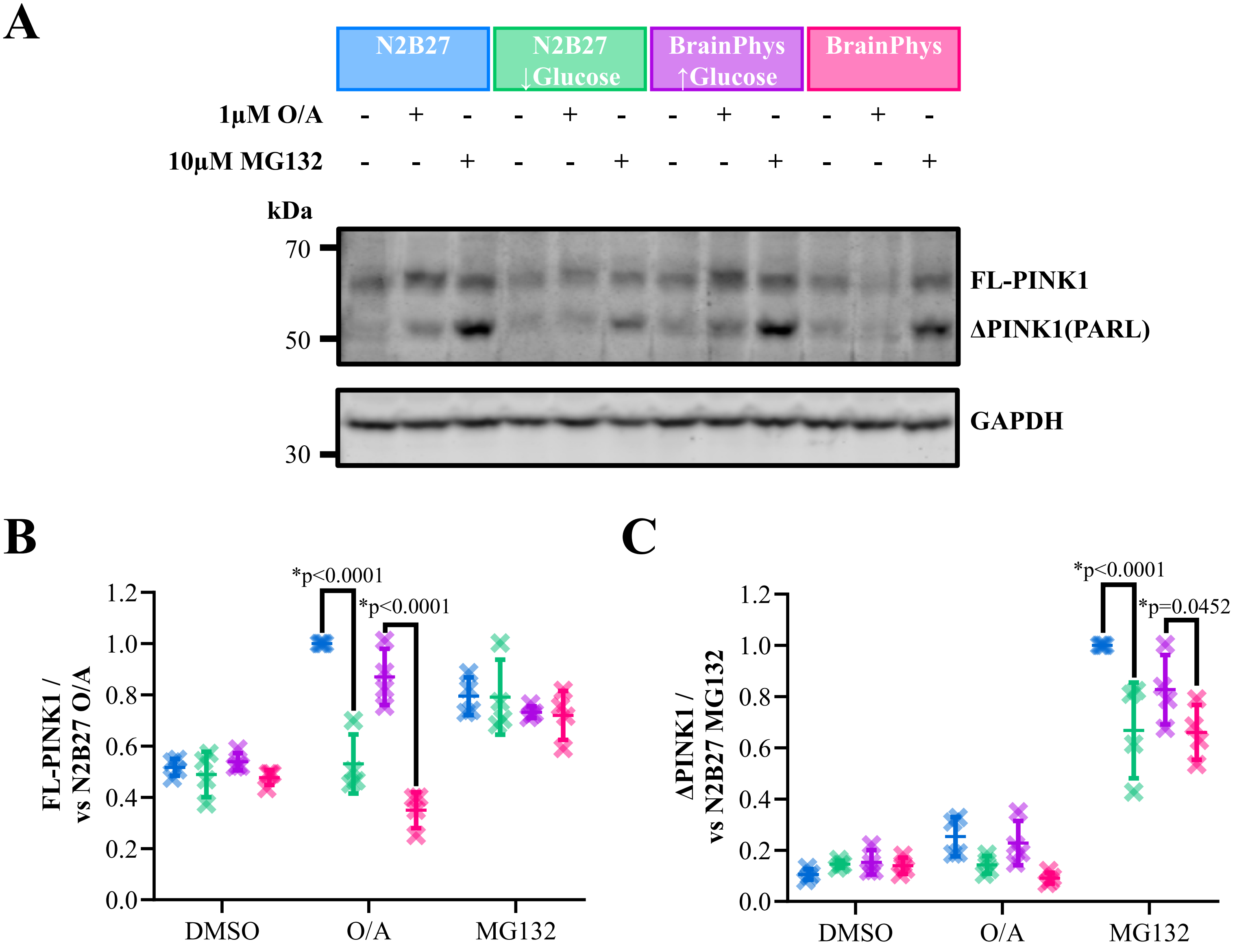
